# *De novo* annotation of lncRNA *HOTAIR* transcripts by long-read RNA capture-seq reveals a differentiation-driven isoform switch

**DOI:** 10.1101/2022.06.17.496514

**Authors:** Evdokiia Potolitsyna, Sarah Hazell Pickering, Ave Tooming-Klunderud, Philippe Collas, Nolwenn Briand

**Affiliations:** Department of Molecular Medicine, Institute of Basic Medical Sciences, Faculty of Medicine, University of Oslo, PO Box 1112 Blindern, 0317 Oslo, Norway; Department of Immunology and Transfusion Medicine, Oslo University Hospital, 0424 Oslo, Norway; Centre for Ecological and Evolutionary Synthesis, Department of Biosciences, University of Oslo, Oslo, Norway

**Keywords:** *HOTAIR*, adipose differentiation, adipose stem cells, capture-seq, long-read sequencing, lncRNA isoform

## Abstract

**Background:** LncRNAs are tissue-specific and emerge as important regulators of various biological processes and as disease biomarkers. *HOTAIR* is a well-established pro-oncogenic lncRNA which has been attributed a variety of functions in cancer and native contexts. However, a lack of an exhaustive, cell type-specific annotation questions whether *HOTAIR* functions are supported by the expression of multiple isoforms.

**Results:** Using a capture long-read sequencing approach, we characterize *HOTAIR* isoforms expressed in human primary adipose stem cells. We identify a highly cell type-specific *HOTAIR* isoform and uncover a shift in the *HOTAIR* isoform balance at differentiation onset. Composition of the *HOTAIR* isoform pool is regulated by distinct promoter usage and is under control of hormonal and nutrient-sensing pathways.

**Conclusion:** Our results highlight the complexity and cell type-specificity of *HOTAIR* isoforms and open perspectives on functional implications of these variants and their balance to key cellular processes.

## Background

Long non-coding RNAs (lncRNAs) are increasingly recognized as major regulators of physiological processes and have emerged as biomarkers for disease diagnosis and prognosis [1]. *HOTAIR* (*HOX* Transcript Antisense RNA) is a long antisense transcript of ~500-2200 base pairs (bp) located within, and extending upstream of, the *HOXC11* gene, in the *HOXC* locus on chromosome 12. *HOTAIR* is mostly studied in cancer models where its overexpression promotes cell migration and metastasis by altering gene expression [2–5]. *HOTAIR* is also the most differentially expressed gene between upper- and lower-body adipose tissue, and its expression is induced during adipose differentiation of gluteofemoral adipose stem cells (hereafter referred to as ASCs) [6, 7]. The function of *HOTAIR* in adipose tissue remains, however, unclear. In contrast to its effect in cancer cell lines [8–11], *HOTAIR* overexpression does not affect adipose progenitors gene expression or proliferation rates [7, 12]. These observations point to different functions and mechanisms of action of *HOTAIR* in cancer vs. mesenchymal progenitor cells.

LncRNAs interact with proteins, DNA and other RNAs to regulate gene expression at multiple levels. *HOTAIR* has been detected in the cytoplasm where it can promote ubiquitin-mediated proteolysis by associating with E3 ubiquitin ligases [13] or function as a microRNA sponge [14]. *HOTAIR* is also found in the nucleus, where it can bind chromatin [13, 15, 16] and act as scaffold for chromatin-modifying complexes through binding to the Polycomb repressor complex PRC2 subunit EZH2 [17], a histone H3K27 methyltransferase, and to LSD1/KDM1A, the H3K4/K9 demethylase of the REST/CoREST complex [18]. Recent evidence demonstrates that HOTAIR-PRC2 binding can be modulated by changes in *HOTAIR* structure mediated by RNA binding protein (RBP)-RNA-lncRNA interactions [19].

LncRNA folding into secondary and tertiary structures dictates their interactome, making their function dependent on structural conservation[20]. While the full-length *HOTAIR* sequence is poorly evolutionarily conserved, folding prediction has identified two well-conserved structures in the 5’ and 3’ ends of *HOTAIR* [21]. The currently reported primary structure of the *HOTAIR* gene is complex, with 2 predicted promoters, multiple predicted transcription start sites (TSSs) and potential splice sites leading to several isoforms [22–25]. *HOTAIR* transcripts can harbor small exon length variations [26], or alternative splice site usage that eliminates the PRC2 [18] or RBP [19] binding domains. Therefore, distinct *HOTAIR* splice variants likely have distinct functions, warranting an isoform-specific annotation in relevant tissues.

Current reference annotations for lncRNAs are incomplete due to their overall low expression level, weak evolutionary conservation, and high tissue specificity [27]. Identification of lncRNA isoforms using shortread RNA sequencing (RNA-seq) is challenging because almost every exon can be alternatively spliced [30], and short reads cannot resolve the connectivity between distant exons. Long-read sequencing technologies can address this challenge by covering the entire RNA sequence in a single read, enabling mapping of isoform changes that may impact lncRNA structure and function [28].

Here, we combine long-read Capture-seq and Illumina short-read RNA-seq to resolve changes in the composition of *HOTAIR* isoforms in a well-characterized adipogenic differentiation system [29]. We uncover a temporal shift in the composition of *HOTAIR* isoforms upon induction of differentiation, regulated by distinct promoter usage and hormonal and nutrient-sensing pathways.

## Results

### *HOTAIR* is highly expressed in ASCs and regulated during adipogenesis

We first assessed *HOTAIR* expression level in ASCs versus cancer cell lines where *HOTAIR* function has been previously studied [30]. We find that *HOTAIR* expression level in ASCs from two unrelated donors is higher or comparable to that of cancer cell lines (**Fig.1a**), confirming the relevance of primary ASCs as a model system to assess the relative abundance of *HOTAIR* isoforms.

**Figure 1.**
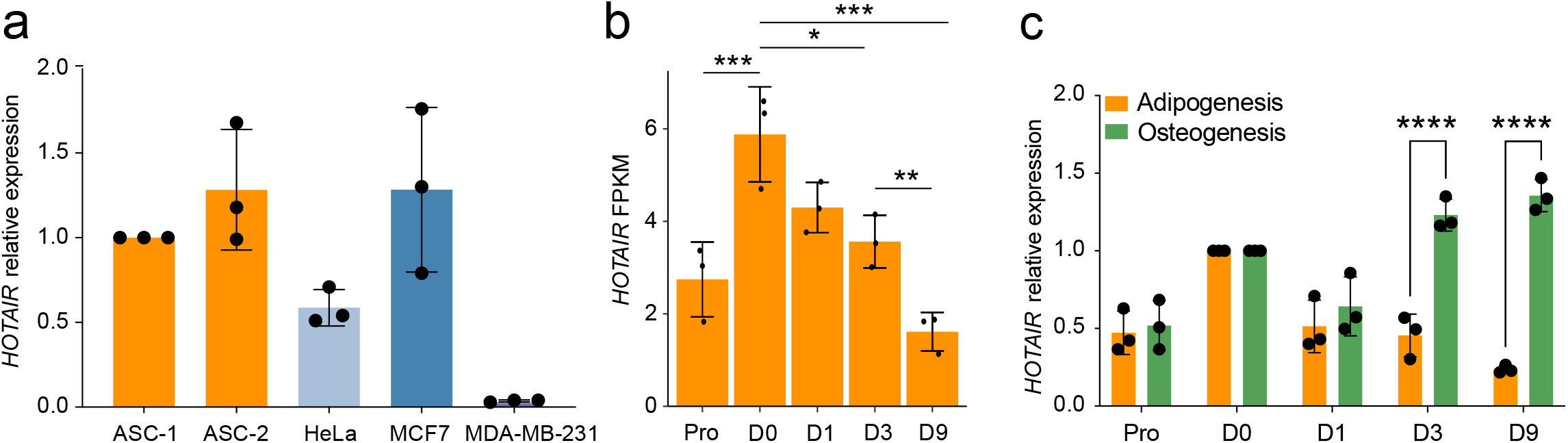
Validation of adipogenesis as a model to study *HOTAIR* isoforms. **a** Quantitative RT-PCR analysis of *HOTAIR* expression in HeLa, MCF7, MDA-MB-231, and ASCs from two independent donors (ASC-1 and ASC-2). **b** Differential expression of *HOTAIR* between time-points analysed by RNA-seq (mean ± SD; *p < 0.05, **p<0.01, ***p<0.001, limma moderated t-statistic; n = 3). **c** Relative *HOTAIR* expression normalized to D0 ASCs during adipose and osteogenic differentiation (mean fold difference ± SD; *p < 0.05, two-way ANOVA; n = 3).

We examined by RNA-seq the transcriptome of ASCs in the proliferating stage (Pro), after cell cycle arrest (day 0; D0), and after 1, 3 and 9 days of adipogenic induction (D1, D3, D9). Hierarchical clustering of differentially expressed genes across time (α < 0.01 between at least two consecutive time points) confirms the upregulation of genes pertaining to the hallmark “Adipogenesis”, including the master adipogenic transcription factor *PPARG* (**Fig. S1**). Moreover, *HOTAIR* displays a biphasic expression profile in this time course, with increased levels upon cell cycle arrest on D0, followed by a progressive downregulation (**Fig. 1b**). *HOTAIR* expression is maintained during osteogenic differentiation (**Fig. 1c; Fig. S2a**), indicating a lineage-specific mode of regulation.

### Identification of main *HOTAIR* isoforms

To identify *HOTAIR* isoforms, we performed PacBio single-molecule, long-read isoform sequencing of captured polyadenylated *HOTAIR* transcripts (PacBio Capture-seq) in proliferating ASCs and during adipose differentiation. Full-length reads were clustered into non-redundant transcripts and aligned to the hg38 genome assembly, providing excellent coverage over the *HOTAIR* locus with sharp exon boundaries compared to short-read Illumina coverage (**Fig. 2a,b**). The Isoseq3 pipeline yielded ~6000 *HOTAIR* isoforms; these were further filtered and merged both across time points and based on internal junctions using Cupcake ToFU [31, 32] or TAMA [33] (**Fig. 2a**), resulting in 34 isoforms (**Fig. 2a,c; Additional file 1, Table S1**).

**Figure 2.**
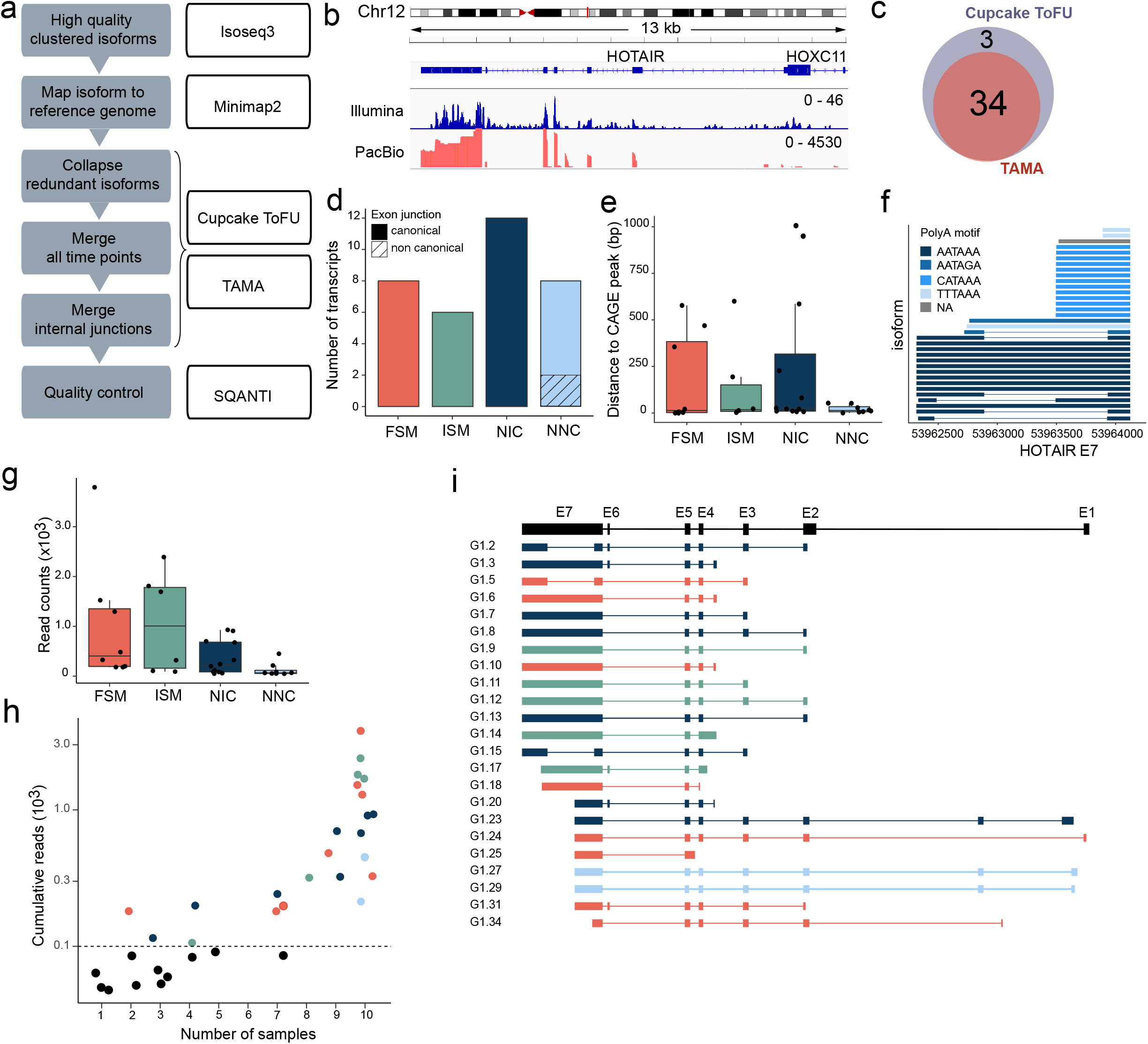
Identification of *HOTAIR* isoforms in differentiating ASCs. **a** Bioinformatic pipeline for identification of HOTAIR isoforms. **b** Representative integrative genomics viewer (IGV) tracks for shortread Illumina RNA-seq (upper) and PacBio capture-seq (lower) coverage tracks on *HOTAIR* (D0). **c** Venn diagram showing the overlap between *HOTAIR* isoforms identified using TAMA and Cupcake ToFU. **d-h** SQANTI characterization of 34 *HOTAIR* transcripts, with: **d** Number of isoforms per SQANTI categories and presence of non-canonical splice sites. **e** Distance in base pairs from the isoform start to the nearest CAGE peak summit from Ref. [43]. **f** HOTAIR exon E7 diagram showing 3’ end polyA tails. **g** Cumulative read counts of the isoforms shown by SQANTI categories. **h** Cumulative read number per isoform vs. number of samples in which the isoform was detected (cutoff: 0.1×10^3^ reads per isoform). **i** Exon structure of 23 high confidence *HOTAIR* isoforms colored by SQANTI categories. FSM: full splice match; ISM: incomplete splice match; NIC: novel in catalog; NNC: novel not in catalog.

To characterize *HOTAIR* isoforms, we first classified transcripts into four main categories using SQANTI. Out of the 34 aforementioned isoforms, we find (i) 8 full splice matches (FSM) transcripts, (ii) 6 incomplete splice matches (ISM) of an annotated (known) transcript, (iii) 12 novel in catalog (NIC) transcripts containing new combinations of annotated splice sites, and (iv) 8 novel not in catalog (NNC) transcripts using at least one unannotated splice site (**Fig. 2d**). Second, we assessed isoform variations at the 5’ and 3’ ends. Transcripts start sites supported by Cap analysis of gene expression (CAGE) data are distributed evenly across SQANTI categories, with 15 transcripts starting within 15 bp of a CAGE peak summit (**Fig. 2e**). Transcripts with a TSS located more than 250 bp away from a CAGE peak likely represent lowly expressed or highly cell type-specific isoforms of *HOTAIR*, explaining the absence of a dedicated CAGE peak [34, 35]. *HOTAIR* full length transcripts also show variable 3’ ends (exon 7; E7) corresponding to the alternative usage of 4 different canonical polyA signals for transcription termination (**Fig. 2f, Additional file 1, Table S2**). Third, we find the highest number of reads for transcripts in FSM, ISM and NIC SQANTI categories (**Fig. 2g**), indicating that most *HOTAIR* transcripts identified here have known exon and splice junction composition.

To identify the top isoforms expressed across differentiation, we further filtered candidates based on read counts (**Fig. 2h**). Only 23 transcripts accumulate more than 100 reads, and those are also detected in at least 2 samples. These 23 high-confidence isoforms arise from multiple TSS usage, alternative splicing and intron retention events, as well as polyA site usage (**Fig. 2i**). Altogether, our PacBio sequencing analysis identifies with high confidence known and novel uncharacterized *HOTAIR* transcript isoforms in our adipose cell system, with notable variation in their TSSs and polyA site usage.

### *HOTAIR* splicing affects LSD1 and PRC2 interacting domains

*HOTAIR* has been described as a scaffold for LSD1 [18] and PRC2 [18], epigenetic modifiers involved in the regulation of adipogenesis [36–38]. The LSD1 binding domain lies in the last 500 bp of *HOTAIR* exon 7 (E7), whereas the PRC2 binding domain spans exons 4 and 5 [39] (**Fig. 3a**).

**Figure 3.**
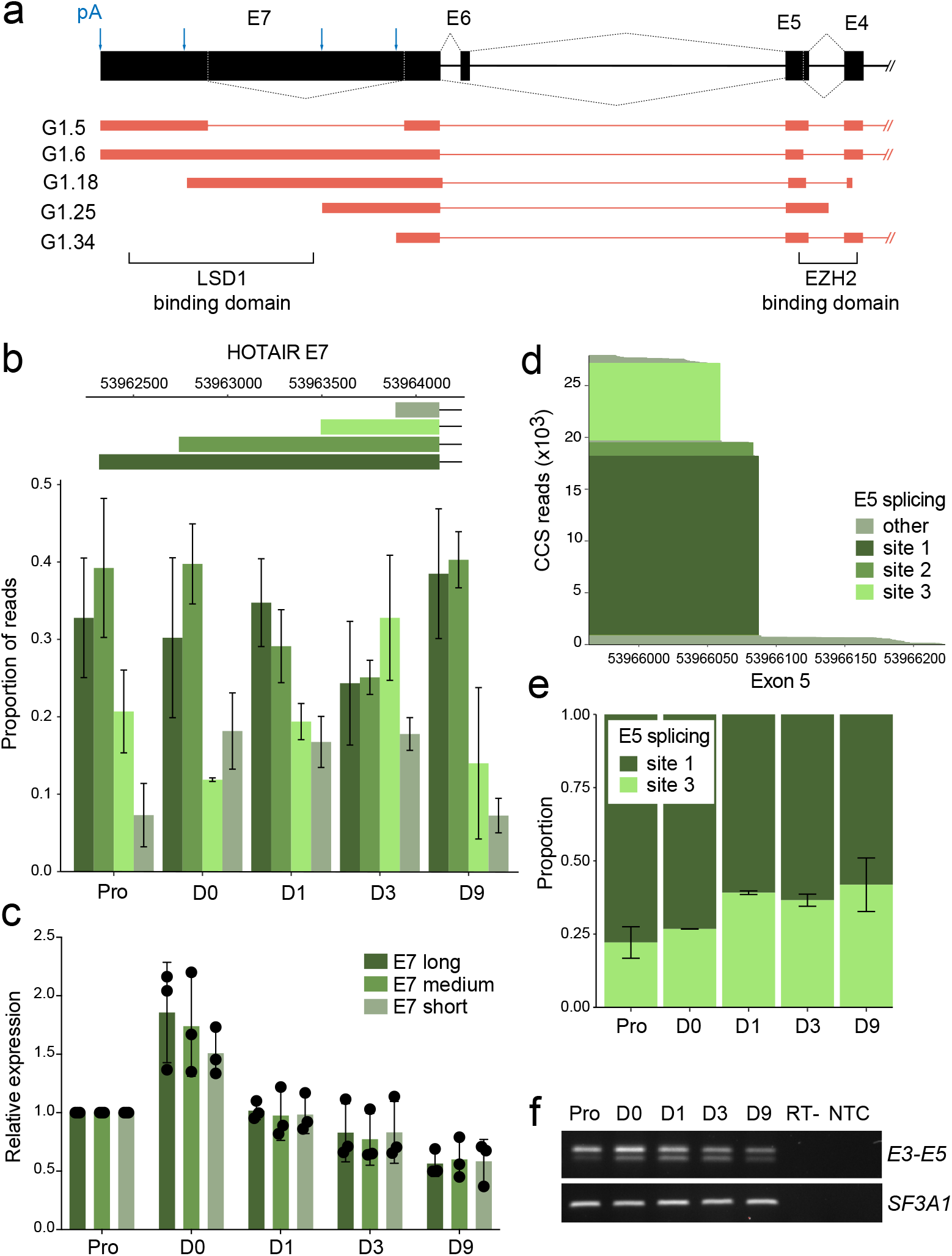
*HOTAIR* splicing across functional domains during adipogenesis. **a** Schematic representation of *HOTAIR* 3’ exons (exons E4 to E7) containing LSD1 and PRC2 binding domains. Alternative splicing events and polyA (pA) sites usage detected in 5 representative *HOTAIR* high confidence isoforms are represented. **b** Proportion of long-read Capture-Seq reads for each E7 length across differentiation time points. **c** qRT-PCR analysis of *HOTAIR* E7 length variation during adipose differentiation. **d** PacBio Capture-Seq analysis of HOTAIR exon E5 alternative splice site usage and **e** of the proportion of major exon E5 splice variants during adipogenic differentiation. **f** Semi-quantitative RT-PCR analysis of *HOTAIR* expression using primers located in exons E3 and E5. *SF3A1* is shown as a loading control. RT-: no reverse transcriptase control; NTC: no template control. Full-length gels are presented in **Additional file 3 Figure S3**.

Inclusion of the LSD1 binding domain in *HOTAIR* isoforms depends on polyA site usage (see **Fig. 3a; Fig. 2f,i**); thus we examined potential changes in exon E7 length during adipogenesis. The proportion of reads for the long forms of E7, harboring the LSD1 binding domain, does not significantly vary during differentiation (**Fig. 3b**), a result confirmed by RT-qPCR (**Fig. 3c**). Hence, variations in *HOTAIR* isoforms during adipogenesis do not in principle impact its LSD1 scaffolding function.

We next investigated alternative splicing events affecting the PRC2 binding domain. Analysis of exon E5 alternative splicing reveals two major alternative splice sites (site 1 and site 3) (**Fig. 3d**). Induction of differentiation promotes splicing at site 3, leading to a slight increase in proportion of the shorter E5 variant (**Fig. 3e**). This splicing event can be readily detected by semi-quantitative RT-PCR analysis of exon E3-E5 expression in ASCs from two independent individuals (**Fig. 3f, Additional file 2, Fig. S2a,b**), emphasizing the robustness of the PacBio approach in capturing relative variations in *HOTAIR* transcript levels during differentiation. Overall, PacBio Capture-seq analysis reveals that *HOTAIR* is expressed as a pool of isoforms with differential ability to bind LSD1 and PRC2. However, adipogenic induction does not significantly affect the proportion of isoform with LSD1 or PRC2 binding capacity.

### Adipogenic induction triggers a switch in *HOTAIR* isoform start sites

One noticeable feature of *HOTAIR* isoform pool is the presence of 9 distinct start sites (**Fig.4a**, see **Fig.2i**). We therefore assessed whether adipogenesis is accompanied by a change in *HOTAIR* TSS usage. We categorized transcripts based on their start exons and evaluated the number of reads for each category in the PacBio Capture-seq dataset (**Fig. 4b**). Only transcripts starting from exons E2, E3, E3.1 and E5 cumulated more than 500 reads during differentiation. Among these, we noted the proportion of isoforms starting from E3 increases after adipogenic induction, whereas E3.1 starting transcripts decrease (**Fig. 4c,d**). RNA-seq analysis confirms this switch in TSS usage, with exon E3 becoming significantly more expressed than E3.1 after differentiation onset (**Fig. 4e**). The sharp drop in E3.1 starting isoforms is readily observed by semi-quantitative RT-PCR using primers spanning exons E3.1-E4, while expression of E3-E5 containing isoforms is maintained during early adipogenesis (**Fig. 4f**, left; **Additional file 2, Fig. S2a,b**). Importantly, osteogenic differentiation of ASCs maintains E3.1 isoforms expression (**Fig. 4d**, right; **Additional file 2, Fig. S2a**), indicating an adipose-specific regulation of *HOTAIR* isoforms. We conclude that adipogenic commitment triggers a switch in *HOTAIR* TSS usage, potentially impacting functional binding domains located in 5’ exons [19].

**Figure 4.**
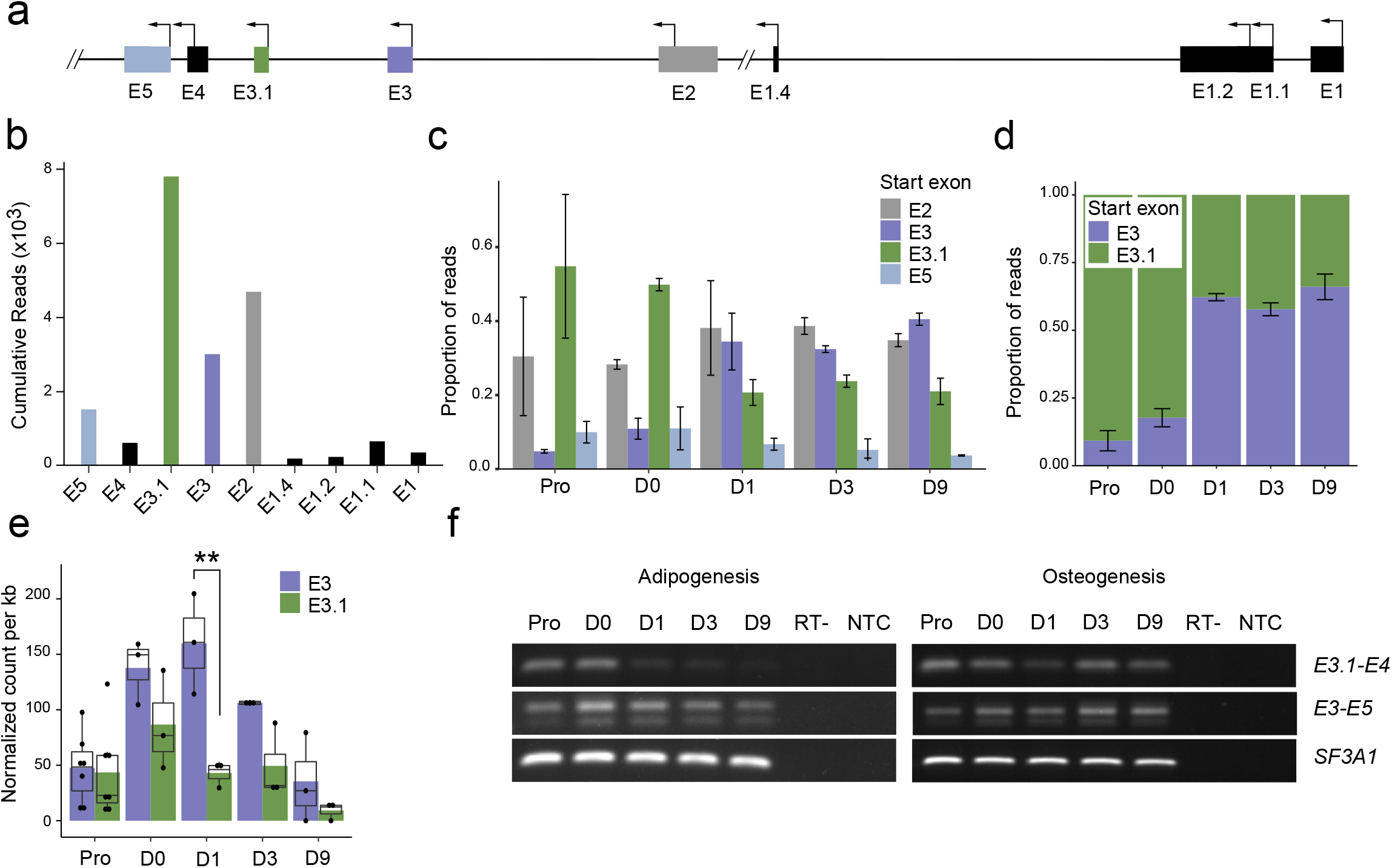
Remodeling of the *HOTAIR* isoform pool during adipogenesis. **a** Schematic representation of *HOTAIR* 5’ exons (exons E1 to E5) showing novel *HOTAIR* exons and 9 alternative TSSs detected by PacBio capture-seq. **b** Cumulative number of PacBio long reads for each *HOTAIR* starting exon. Proportion of read coverage for each start exon quantified from long-read Capture-Seq data for **c** *HOTAIR* isoforms cumulating > 500 reads over the time course and **d** E3- and E3.1-starting isoforms. **e** Normalized RNA-seq read count for E3 and E3.1 exons during differentiation (n = 3, paired t-test) **f** Semi-quantitative RT-PCR analysis of *HOTAIR* expression using primers located in *HOTAIR* exons E3.1-E4 and E3-E5. *SF3A1* is shown as a loading control. A representative image from one of three independent experiments is shown for adipogenesis (left) and osteogenesis (right); see also **Additional file 2 Fig. S2a**. RT-: no reverse transcriptase control; NTC: no template control. Full-length gels are presented in **Additional file 3 Fig.S5.**

### ASCs express a cell type-specific *HOTAIR* isoform

To assess cell type- and tissue-specificity of E3.1 starting *HOTAIR* isoforms, we used Snaptron, a search engine for querying splicing patterns in publicly available RNA-seq datasets [45]. We found only 24 cell samples with more than 10 reads containing the E3.1-E4 junction, while the E3-E4 junction was detected in 624 samples (**Fig. 5a**; **Additional file 1**, **Table S3**). We confirmed by semi-quantitative RT-PCR the expression of exon E3.1 in cultured primary myoblasts, BJ fibroblasts and HEK 293T cells – albeit to lower levels than in ASCs – and its absence in HeLa cells (**Fig. 5b**).

**Figure 5.**
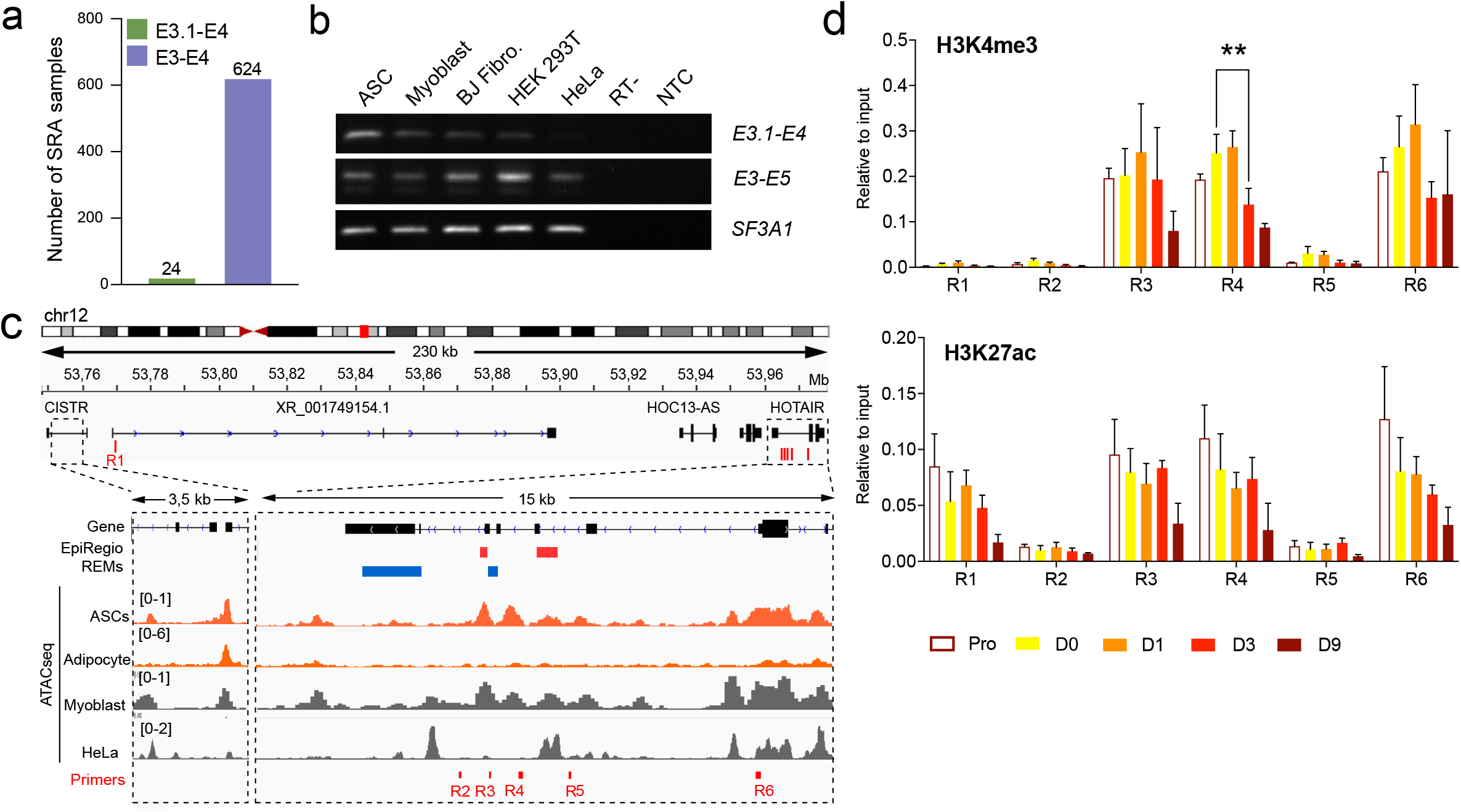
*HOTAIR* isoform expression is highly cell-type specific. **a** Number of SRA samples with 10 or more junction spanning reads for exons E3.1-E4 and E3-E4. **b** Semi-quantitative RT-PCR analysis of *HOTAIR* expression in ASCs, primary myoblasts, BJ fibroblasts, HEK293T and HeLa cells using primers located in *HOTAIR* exons E3.1-E4 and E3-E5. *SF3A1* is shown as a loading control. Full-length gels are presented in **Additional file 3 Fig.S6. c** IGV tracks over *HOTAIR* locus showing predicted HOTAIR regulatory elements (REMs) from the EpiRegio database [44, 60] (red: activating, blue: repressing), average ATAC-Seq tracks from ASCs [40, 41], Myoblasts [42] and HeLa cells [43], and ChIP primers location. **d** ChIP-qPCR analysis of histone modifications during adipogenesis of ASCs (mean ± SEM of n ≥ 3 independent differentiation experiments; **p < 0.01 Mixed effect analysis).

To gain insight into the regulation of *HOTAIR* isoform expression, we examined the chromatin accessibility landscape of the *HOTAIR* locus in published Assay for Transposable Accessible Chromatin (ATAC)-seq data [40–43] (**Fig. 5c**). We find two regions of accessible (‘open’) chromatin in ASCs (R3, R4), which our chromatin immunoprecipitation (ChIP)-seq data show are also enriched in H3K4me3 and H3K27ac, histone modifications characterizing active regulatory sites (**Fig. 5c, d**). Region R3 coincides with both activating and repressing regulatory element (REM) annotations by EpiRegio [44], suggesting it constitutes a bivalent regulator for *HOTAIR* or nearby genes expression. Region R4, located immediately upstream of exon E3.1 is also in an ‘open’ chromatin in myoblasts but not in HeLa cells, which respectively do and do not express E3.1-starting transcripts (**Fig. 5b,c**), and likely represents the active promoter for E3.1-starting isoforms. In contrast, these regions are in a ‘closed’ configuration in mature adipocytes, consistent with *HOTAIR* downregulation during adipogenesis (see **Fig.1b**). Thus, *HOTAIR* displays active regulatory sites in ASCs, in line with the expression of specific isoforms. However, *HOTAIR* upregulation upon cell cycle arrest is not accompanied by changes of H3K4me3 and H3K27ac levels, and loss of H3K4me3 at region 4 occurs only at later differentiation time points (D3 and D9). Thus, modulation of *HOTAIR* levels during adipogenesis is not mediated by epigenetic regulation (**Fig. 5d**).

### *HOTAIR* stability increases upon cell cycle arrest

*HOTAIR* half-life varies according to cell type, from 4 h in HeLa cells [13] to more than 7 h in primary trophoblast cells [45]. We therefore asked whether increased *HOTAIR* levels upon cell cycle arrest (D0; see **Fig. 1b**) could relate to a change in lncRNA stability. Treatment with Flavopiridol to inhibit RNA polymerase II (Pol II) transcription reveals an increase in *HOTAIR* half-life from 3 h in proliferating ASCs to > 4 h in D0 cells (**Fig. 6a**), while cell cycle arrest does not impact the stability of control short-lived *CEBPD* or long-lived *GAPDH* mRNA (**Fig. 6b,c**). Thus, while ATAC-seq data indicate that the *HOTAIR* isoform balance is likely regulated by cell type-specific transcription factors, the global increase in *HOTAIR* levels observed at D0 results at least in part from an increase in its stability.

**Figure 6.**
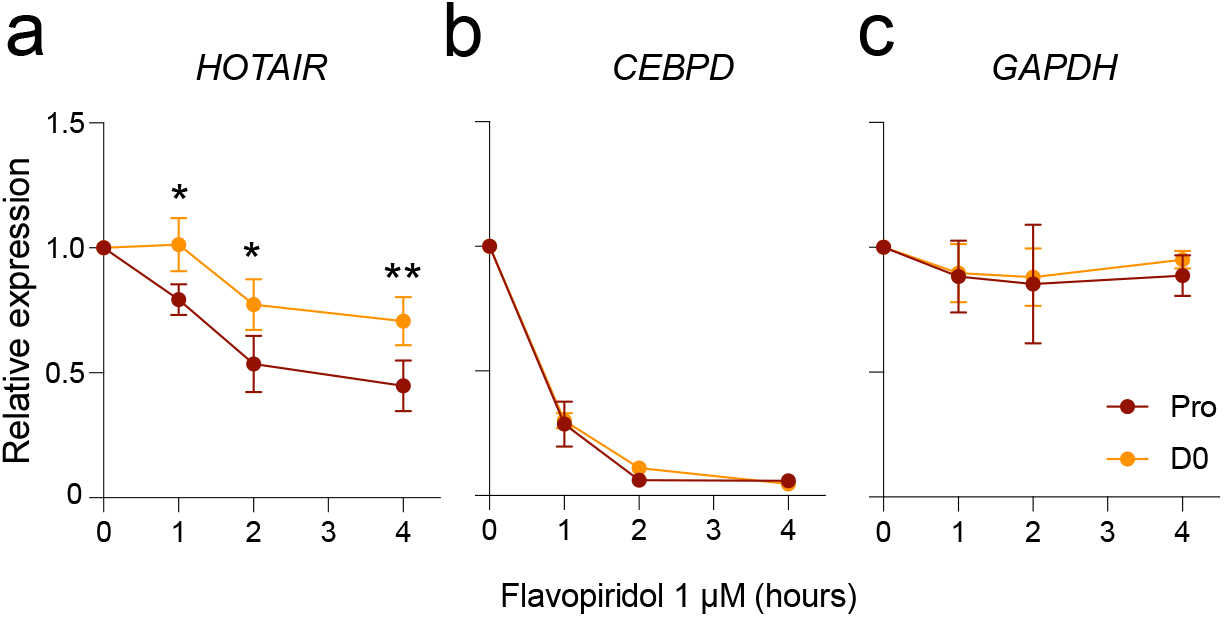
*HOTAIR* stability increases upon growth arrest. RT-qPCR analysis of **a** *HOTAIR*, **b** *CEBPD* and **c** *GAPDH* levels after flavopiridol treatment (mean ± SD; *p < 0.05, **p < 0.01, two-way ANOVA; n = 3 independent experiments).

### Glucocorticoids and cAMP levels differentially regulate *HOTAIR* isoforms

Finally, we asked whether the sharp decrease in E3.1 starting isoforms observed upon differentiation induction correlates with ASC engagement into an adipogenic program, or rather with a direct effect of cytokines used in the differentiation cocktail. To test this, we induced D0 ASCs for 24 h with the full adipogenic cocktail (D1) or with each cytokine used in the cocktail separately: insulin, acting through the insulin receptors; the phosphodiesterase inhibitor IBMX, increasing intracellular cAMP levels; dexamethasone, acting through the glucocorticoid receptor; indomethacin, inducing C/EBPβ expression. We confirm that only the full adipogenic cocktail elicits *PPARG* expression and thus adipogenic commitment (**Fig. 7a,** left panel). While insulin or indomethacin alone do not alter *HOTAIR* expression relative to unstimulated D0 cells (all isoforms; **Fig. 7a,** right panel), IBMX represses both E3.1 and E3 containing isoforms (**Fig. 7b**; **Additional file 2, Fig. S2c**), resulting in a significant reduction of *HOTAIR* expression compared to D0 (**Fig. 7a**). Intriguingly, dexamethasone alone seems to reduce E3.1-starting isoform expression (**Fig. 7b**, **Additional file 2, Fig. S2c**), while provoking a slight increase in expression of E3-containing isoforms; this could account for the maintenance of total *HOTAIR* level detected by RT-qPCR (**Fig. 7a**). Collectively, our data show that during adipogenic differentiation, the balance of *HOTAIR* isoforms is controlled by major metabolic regulators such as glucocorticoids and cAMP levels.

**Figure 7.**
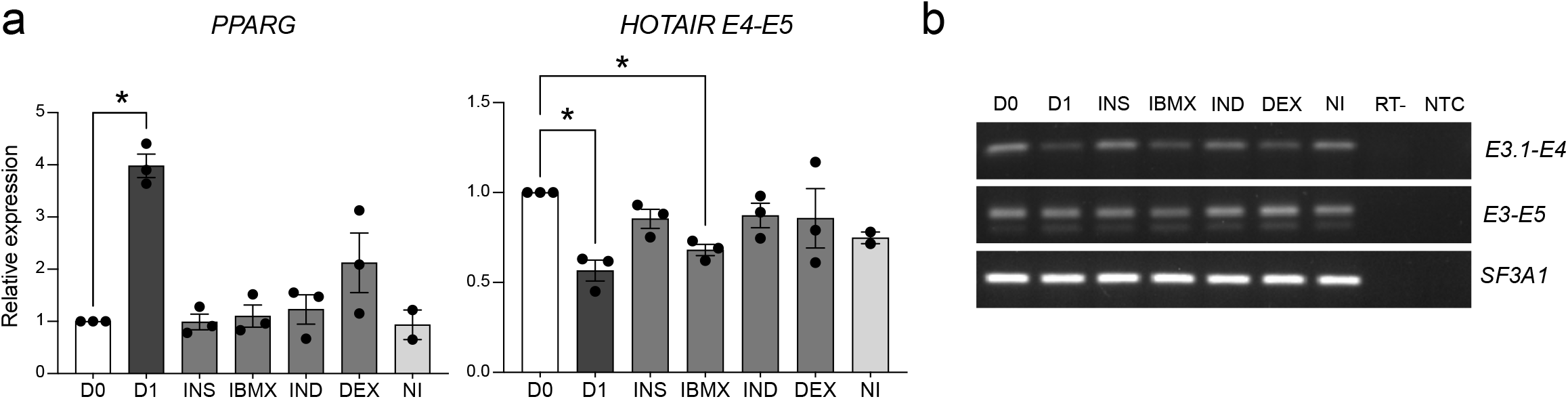
Glucocorticoids and cAMP levels differentially regulate *HOTAIR* isoforms. **a** Relative expression of *PPARG* and *HOTAIR* (all isoforms) (mean fold difference ± SD; *p < 0.05, two-way ANOVA; n = 3) and **b** semi-quantitative RT-PCR analysis of *HOTAIR* expression using primers located in *HOTAIR* exons E3.1-E4 and E3-E5 in growth arrested ASC (D0) treated for 24h with a full adipogenic cocktail (D1) or insulin (INS), 3-Isobutyl-1-methylxanthine (IBMX), dexamethasone (DEX) and indomethacine (IND) only. NI: non-induced; RT-: no reverse transcriptase control; NTC: no template control. A representative image from one of three independent experiments is shown; see also **Additional file 2 Fig. S2c**. Full-length gels are presented in **Additional file 3 Fig.S7.**

## Discussion

LncRNAs can modulate important biological and pathophysiological processes through their interaction with multiple partners. Our long-read Capture-seq of *HOTAIR* unveils a complex and dynamic pool of isoforms in differentiating ASCs. Adipogenic induction triggers a switch in the expression of main *HOTAIR* isoforms which likely impacts on its structure and interactome. Our results emphasize the importance of robust lncRNA annotation in the tissue of interest prior to functional characterization.

A single lncRNA locus can generate many transcript variants through alternative TSS usage, polyadenylation sites and splicing events [46]. We find that a previously undescribed *HOTAIR* variant constitutes the major isoform in human adipose stem cells, and show that not only lncRNA expression level, but also the isoform pool expressed can be highly cell type-specific, adding another layer of complexity to the regulation of biological processes by lncRNAs.

The chromatin landscape of the *HOTAIR* locus in ASCs is consistent with a promoter upstream of *HOTAIR* exon E3.1, which is in an ‘open’ and active epigenetic configuration in ASCs and myoblasts, where *HOTAIR* E3.1 starting isoforms are expressed, but not in HeLa cells. However, adipogenic induction results in only mild and delayed changes in active histone modification at this site, suggesting that expression of various *HOTAIR* isoforms is rather regulated by differentiation-stage specific transcription factors. In line, alternative promoter usage is elicited downstream of adipogenic, but not osteogenic, signaling pathways. Additionally, increased *HOTAIR* levels upon cell cycle arrest (D0 cells) also correlates with an increase in its stability. Interestingly, interaction with the RNA-binding protein HuR, a negative regulator of adipogenesis [47], reduces *HOTAIR* stability [13]. Alternatively, increased *HOTAIR* stability could result from its binding to a cell stage-specific protein partner. Collectively, our results indicate a tight, multifactorial regulation of isoform pool during adipogenesis and support the idea of the functional importance of the isoform switch.

In cancer cell lines, *HOTAIR* has been reported to scaffold for chromatin regulators with broad effects on gene expression [48–50]. Our long-read sequencing data reveal multiple polyA site usage leading to the exclusion of the LSD1 binding domain at the 3’ end, or alternative splicing events affecting the PRC2 minimal binding domain. However, the proportion of *HOTAIR* isoforms containing LSD1 or PRC2 domains does not vary during adipose differentiation, suggesting that *HOTAIR*’s role in adipogenesis is independent from its epigenetic scaffold function. Supporting this idea, transient *HOTAIR* depletion does not significantly affect gene expression in bone marrow-derived adipose progenitors [12].

Recent studies have confirmed that a non-protein-coding locus can give rise to functionally distinct transcript isoforms [46, 51–53]. Of note, the switch in *HOTAIR* start site upon differentiation induction leads to the inclusion of *HOTAIR* exon 3 containing a protein binding domain [19], which likely alters *HOTAIR* function. Another intriguing possibility is that short sequence variations at the 5’ end impact *HOTAIR*’s secondary structure and thus the folding of functional domains. Observation of *HOTAIR* structure using atomic force microscopy reveals multiple dynamic conformations [54]. It is therefore conceivable that *HOTAIR* functions via conformational changes, induced by or resulting from interactions with protein partners. Hence, variations in isoform composition likely results in cell type specific structures and interactomes, providing a rational for the divergent roles of *HOTAIR* in primary cells and cancer cell lines.

## Conclusions

We generate the first cell type-specific, comprehensive catalog of *HOTAIR* isoforms in a physiological context and describe novel *HOTAIR* isoforms, alternative splicing events, and multiple start site usage. We uncover a shift in the *HOTAIR* isoform balance during early adipogenesis which is regulated by major metabolic pathways. The variability of *HOTAIR* isoforms opens new perspectives for studies in (patho)physiological contexts.

## Methods

All methods were performed in accordance with the guidelines and regulations of the University of Oslo.

### Cell culture and differentiation

ASCs from two non-obese donors were cultured in DMEM/F12 with 10% fetal calf serum and 20 ng/ml basic fibroblast growth factor (Pro). Upon confluency, growth factor was removed, and cells were cultured for 72 h before induction of differentiation (D0). For adipose differentiation, ASCs were induced with 0.5 μM 1-methyl-3 isobutyl xanthine, 1 μM dexamethasone, 10 μg/ ml insulin and 200 μM indomethacin. For osteogenic differentiation, ASCs induced with 0.1 μM dexamethasone, 10 mM β-glycerophosphate and 0.05 mM L-ascorbic acid-2 phosphate. Differentiation media was renewed every 3 days, and samples were harvested on D1, D3 and D9 after induction. Differentiation experiments were done in at least biological triplicates. HeLa cells (American Type Culture Collection; CCL-2) were cultured in MEM medium containing Glutamax (Gibco), 1% non-essential amino acids and 10% fetal calf serum. MDA-MB-231 and MCF-7 cells were cultured in DMEM containing 10% fetal calf serum. Human myoblasts were cultured as described [55]. BJ fibroblasts were cultured in DMEM/F12 with 10% fetal calf serum and 20 ng/ml basic fibroblast growth factor. HEK293T were cultured in DMEM/F12 with 10% fetal calf serum.

### RT-qPCR and semi-quantitative PCR

Total RNA was isolated using RNeasy kit (QIAGEN) and 1 μg was used for cDNA synthesis using the High-Capacity cDNA Reverse Transcription Kit (ThermoFisher). RT-PCR was done using IQ SYBR green (Biorad) with *SF3A1* as reference gene. PCR conditions were 95°C for 3 min and 40 cycles of 95°C for 30 s, 60°C for 30 s, and 72°C for 20 s. Semi-quantitative PCR was done using a PCR Master Mix (ThermoFisher) with the following conditions: 95°C for 3 min and 30 cycles of 95°C for 30 s, 60°C for 30 s, and 72°C for 30 s. Products were separated in a 2.5% agarose gel with Tris-Borate-EDTA buffer. PCR primers are listed in **Additional file 1 Table S4.** Uncropped gels are presented in **Additional file 3, Figure S3 to S7**.

### PacBio Capture-Seq

RNA samples from 2 differentiation time courses with 5 time points were used to synthesize full-length barcoded cDNA libraries using the Template Switching RT Enzyme Mix (NEB). Libraries were prepared using Pacific Biosciences protocol for cDNA Sequence Capture Using IDT xGen® Lockdown® probes (https://eu.idtdna.com/site/order/ngs). A pool of 100 probes against all known *HOTAIR* isoform sequences was designed using the IDT web tool. Full-length cDNA was cleaned up using Pronex beads, and 1200 ng was used for each hybridization reactions. Library was sequenced in one 8M SMRT cell on a Sequel II instrument using Sequel II Binding kit 2.1 and Sequencing chemistry v2.0.

### Transcript identification from targeted long-read sequencing

CCS sequences were generated for the entire dataset using the Circular Consensus Sequence pipeline (SMRT Tools v 8.0.0.80502) with minimum number of passes 3 and minimum accuracy 0.99. CCS reads were demultiplexed using the Barcoding pipeline (SMRT Tools v7.0.0.63823). Iso-Seq analysis was performed using the Iso-Seq pipeline (SMRT Link v7.0.0.63985) using default settings. Only clustered isoforms with at least 2 subreads, 0.99 quality score and containing a polyA tail of at least 20 base pairs (bp) were used. Isoforms were aligned to the hg38 genome with minimap2 v.2.17 [56]. Primary alignments to the *HOTAIR* locus (chr12: 53,962,308-53,974,956) were selected as target *HOTAIR* transcripts if they had a mapping quality above 20 and less than 50 clipped nucleotides (samclip; https://github.com/tseemann/samclip). Single exon and sense transcripts were also filtered.

*HOTAIR* transcripts were collapsed with both Cupcake Tofu [31] (https://github.com/Magdoll/cDNA_Cupcake) and TAMA [33] (https://github.com/GenomeRIK/tama). Parameters --dun-merge-5-shorter and –x capped were used for cupcake and TAMA respectively, to prevent shorter transcript models from being merged into longer ones. For TAMA, –z 100 was also set to increase the allowed 3’ variability. To combine transcript lists between timepoints, collapsed transcripts with at least 50 full length reads within one sample were merged using cupcake chain_samples.py or tama_merge.py. Initially this produced ~80 HOTAIR transcripts, with much of the variation in the ends of 3’ and 5’ exons. To achieve a final transcript list, collapsed transcripts were merged on internal junctions only by increasing the allowed 5’ and 3’ variability (with TAMA options -a 300 -z 2000) and ranking libraries by the number of polyadenylated *HOTAIR* reads. This resulted in 34 TAMA isoforms.

Final cross-validation was conducted by running SQANTI with unclustered ccs reads as the “novel long read-defined transcriptome” and the top 34 TAMA isoforms as the “reference annotation”. This assigned full length reads to their corresponding TAMA isoform and reads annotated as full-splice match were counted for each isoform. SQANTI was used again to classify transcripts against existing reference annotations and to search for polyA motifs near transcript ends from a list of potential human motifs (**Additional file 1 Tables S2 and S3**) [57] (https://github.com/ConesaLab/SQANTI).

### Snaptron

We searched Snaptron’s SRA V2 database, which contains ~49000 public samples, for experiments with reads overlapping *HOTAIR* exon-exon junctions with the web query http://snaptron.cs.jhu.edu/srav2/snaptron?regions=HOTAIR&rfilter=annotated:1. Sample IDs for experiments with at least 10 exon spanning reads were extracted and cell type information accessed via a matching metadata query. A simplified version of the exon E3.1 metadata table is presented in **Additional file 1 Table S3**.

### Chromatin immunoprecipitation

Cells (2×10^6^/ChIP) were cross-linked with 1% formaldehyde for 10 min and cross-linking was stopped with 125 mM glycine. Cells were lysed for 10 min in ChIP lysis buffer (1% SDS, 10 mM EDTA, 50 mM Tris-HCl, pH 8.0, proteinase inhibitors, 1 mM PMSF, 20 mM Na Butyrate) and sonicated for 30 sec ON/OFF for 10 min in a Bioruptor®Pico (Diagenode) to generate 200-500 bp DNA fragments. After sedimentation at 10,000 g for 10 min, the supernatant was collected and diluted 5 times in RIPA buffer (140 mM NaCl, 10 mM Tris-HCl pH 8.0, 1 mM EDTA, 0.5 mM EGTA, 1% Triton X-100, 0.1% SDS, 0.1% sodium deoxycholate, protease inhibitors, 1 mM PMSF, 20 mM Na Butyrate). After a 100 μL sample was removed (input), diluted chromatin was incubated for 2 h with antibodies (2.5 μg/100 μL) coupled to magnetic Dynabeads Protein A (Invitrogen). ChIP samples were washed 4 times in ice-cold RIPA buffer, crosslinks were reversed and DNA was eluted for 2h at 68°C in 50 mM NaCl, 20 mM Tris-HCl pH 7.5, 5 mM EDTA, 1% SDS and 50 ng/μl Proteinase K. DNA was purified and dissolved in H2O. ChIP DNA was used as template for quantitative (q)PCR using SYBR® Green (BioRad), with 95°C denaturation for 3 min and 40 cycles of 95 °C for 30 sec, 60 °C for 30 sec, and 72 °C for 30 sec. Primers used for ChIP are listed in **Additional file 1**, **Table S4**.

### Statistical analyses

Statistics were performed with GraphPad Prism 9.2.0 (https://www.graphpad.com/).

## Abbreviations

ASCs: adipose stem cells
ChIP: chromatin immunoprecipitation
GSAT: gluteofemoral subcutaneous adipose tissue
HOTAIR: HOX Transcript Antisense RNA
LncRNA: long non-coding RNA
PRC2: polycomb repressor complex 2

## Declarations

### Ethics approval and consent to participate

The study was approved by the Regional Committee for Research Ethics for Southern Norway (REK 2013/2102 and REK 2018-660), and all subjects gave written informed consent.

### Consent for publication

Not applicable.

### Availability of data and material

The dataset supporting the conclusions of this article is available in the SRA repository under accession PRJNA730802 (https://www.ncbi.nlm.nih.gov/bioproject/PRJNA235292) and the filtered transcript list was submitted to GenBank (see accession numbers in **Additional file 1, Table S1**).

ATAC-seq data were obtained from GEO accession GSE118500 (https://www.ncbi.nlm.nih.gov/geo/query/acc.cgi?acc=GSE118500), GSE139571 (https://www.ncbi.nlm.nih.gov/geo/query/acc.cgi?acc=GSE139571) and GSE157399 (https://www.ncbi.nlm.nih.gov/geo/query/acc.cgi?acc=GSE157399).

CAGE clusters and transcripts from [34] were obtained from FANTOM (https://fantom.gsc.riken.jp/5/suppl/Hon_et_al_2016/data/assembly/lv3_robust/). Other HOTAIR transcripts were downloaded from ensembl v95 [58] (http://jan2019.archive.ensembl.org/index.html) or from the UCSC browser at [59].

Regulatory elements (REMs) [44] associated with the *HOTAIR* locus were queried from the EpiRegio database (https://epiregio.de/geneQuery/; accessed: 23/04/2021).

### Authors’ contributions

EP and NB generated data. SHP performed bioinformatic analysis. ATK performed *HOTAIR* capture and long-read sequencing. EP, SHP and NB interpreted data and made figures. NB supervised the work. EP, SHP, PC and NB wrote or revised the manuscript. All authors read and approved the final manuscript.

## Acknowledgements

We thank Anita Løvstad Sørensen for technical assistance and the Norwegian Sequencing Center (Oslo University Hospital) for professional services.

## Supplementary Information

**Additional file 1** (Excel file)

**Table S1**: SQANTI characterization of *HOTAIR* isoforms identified with TAMA.

**Table S2**: List of human polyA motifs for SQANTI analysis

**Table S3**: Snaptron summary of RNA sequencing data containing *HOTAIR E3.1-E4* splice junction.

**Table S4**: List of primers used in this study.

**Additional file 2** (pdf)

**Figure S1**. Validation of adipogenic differentiation efficiency.

**Figure S2**. Semi-quantitative RT-PCR replicates.

**Additional file 3** (pdf)

**Figure S3.** Uncropped gels for **Fig.3f** and **Additional file 2, FigS2a** left panel

**Figure S4.** Uncropped gels for **Additional file 2, Fig. S2b**

**Figure S5.** Uncropped gels for **Fig.4f** and **Additional file 2, Fig. S2a** right panel

**Figure S6.** Uncropped gels for **Fig.5b**

**Figure S7.** Uncropped gels for **Fig.7b** and **Additional file 2 FigS2c**

### Additional file 2

**Figure S1.**
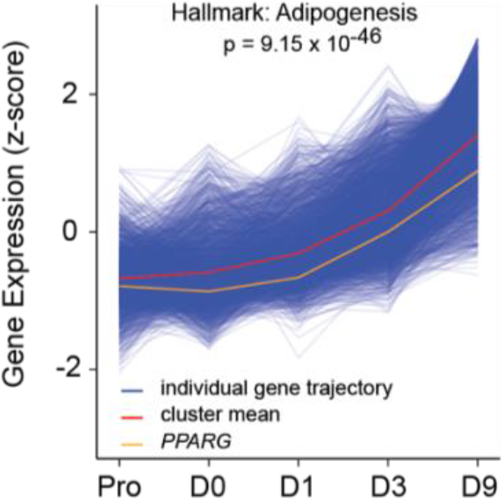
Validation of adipogenic differentiation efficiency. **a** Cluster of differentially expressed genes induced over the adipogenic RNA-seq time-course (n=3, adjusted p-value).

**Figure S2.**
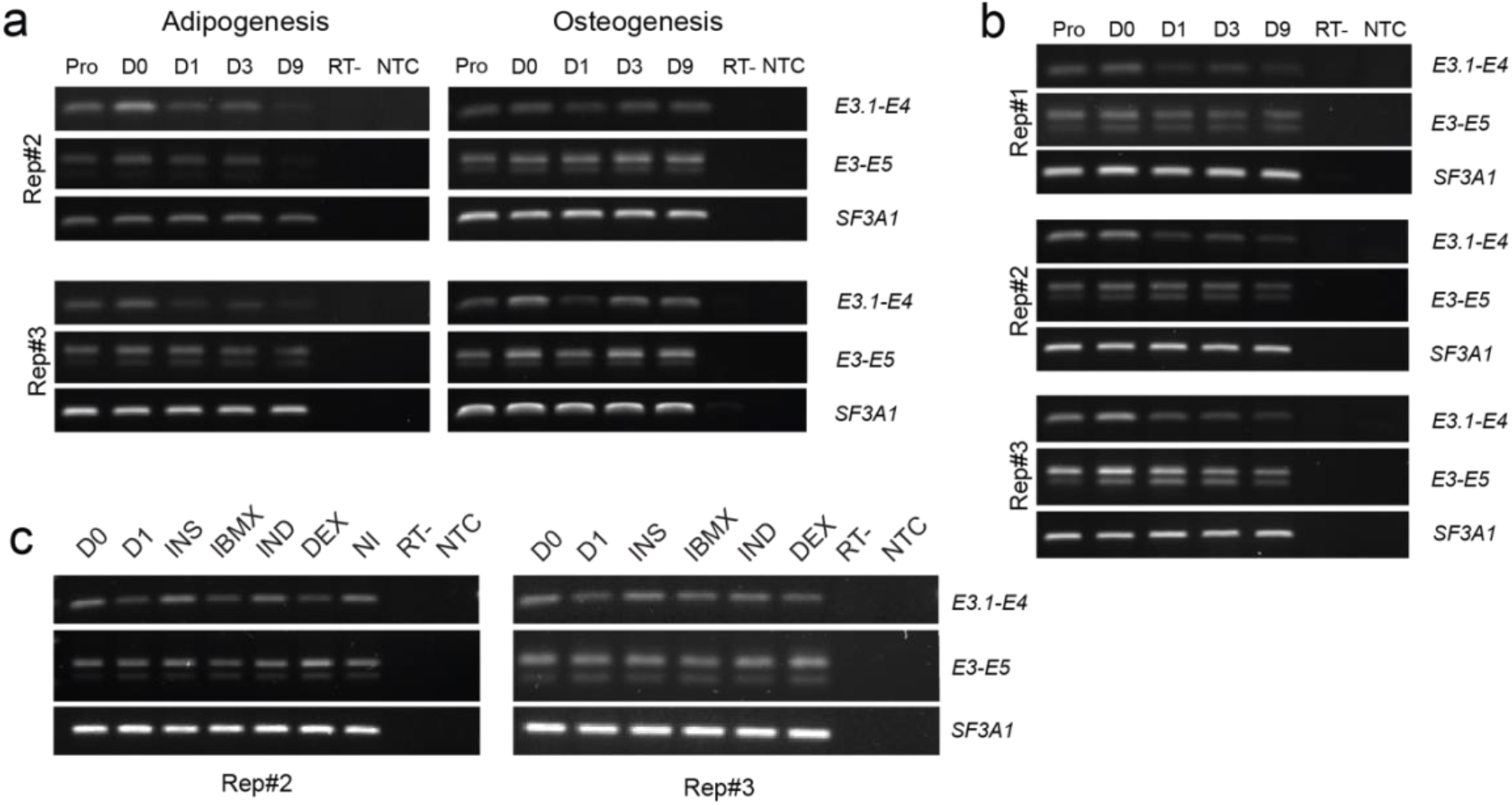
Semi-quantitative RT-PCR replicates. **a** Semi-quantitative RT-PCR analysis of *HOTAIR* isoform expression during adipogenic and osteogenic differentiation using primers located in *HOTAIR* exons E3.1-E4 and E3-E5 in ASCs from donor 1 (replicates #2 and #3; Full-length gels are presented in **Additional file 3 Fig.S1** and **S3**) and **b** donor 2 (Full-length gels are presented in **Additional file 3 Fig.S2**) **c** Semi-quantitative RT-PCR analysis of *HOTAIR* expression using primers located in *HOTAIR* exons E3.1-E4 and E3-E5 in growth arrested ASCs (D0) treated for 24 h with a full adipogenic cocktail (D1) or insulin (INS), 3-Isobutyl-1-methylxanthine (IBMX), dexamethasone (DEX) and indomethacine (IND) only (donor 1; replicates #2 and #3). Full-length gels are presented in **Additional file 3 Fig.S5**. NI: non-induced; RT-: no reverse transcriptase control; NTC: no template control. SF3A1 is shown as a loading control.

### Additional file 3 (pdf)

**Figure S3.**
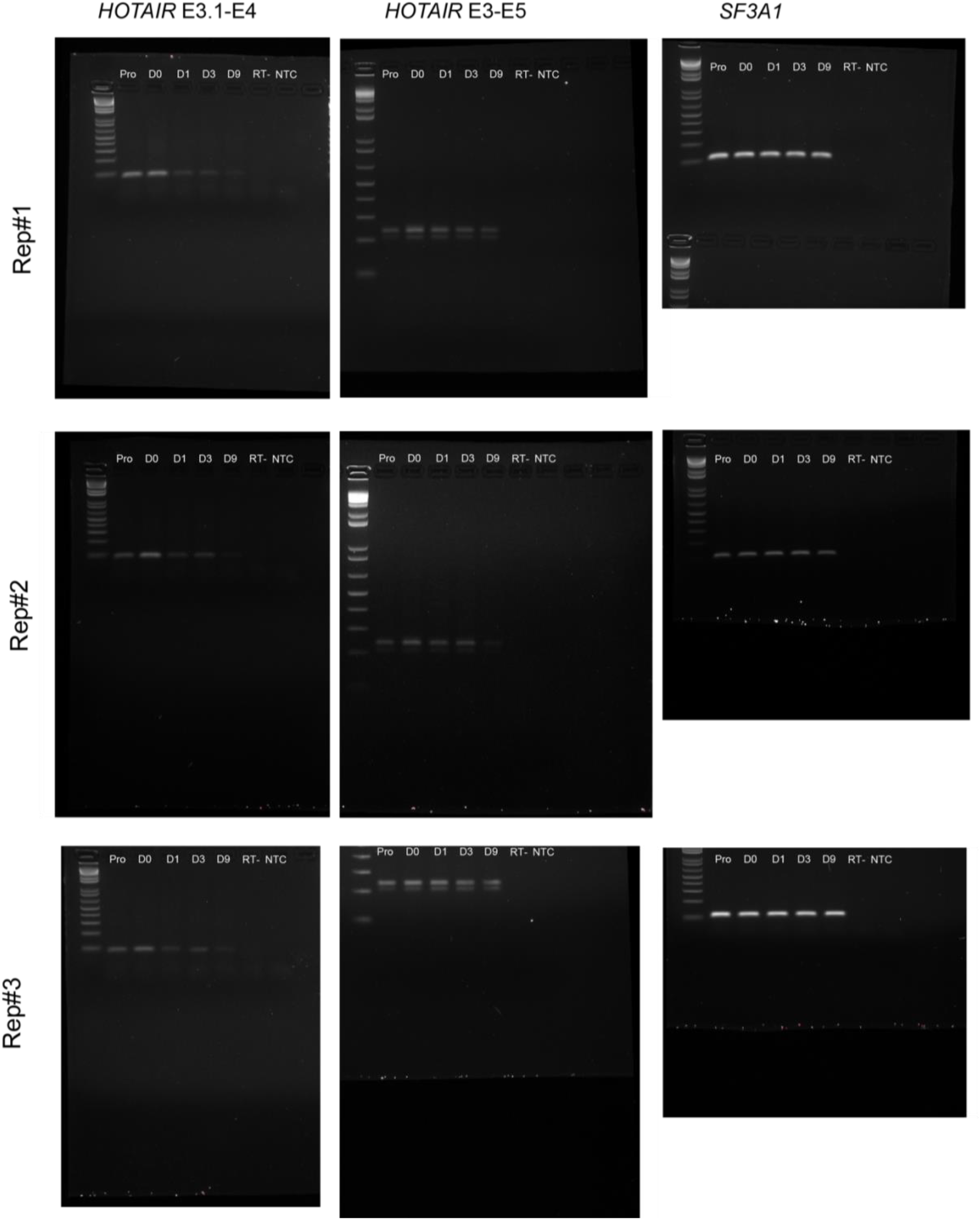
Uncropped gels for **Fig.3f** and **Additional file 2, FigS2a** left panel

**Figure S4.**
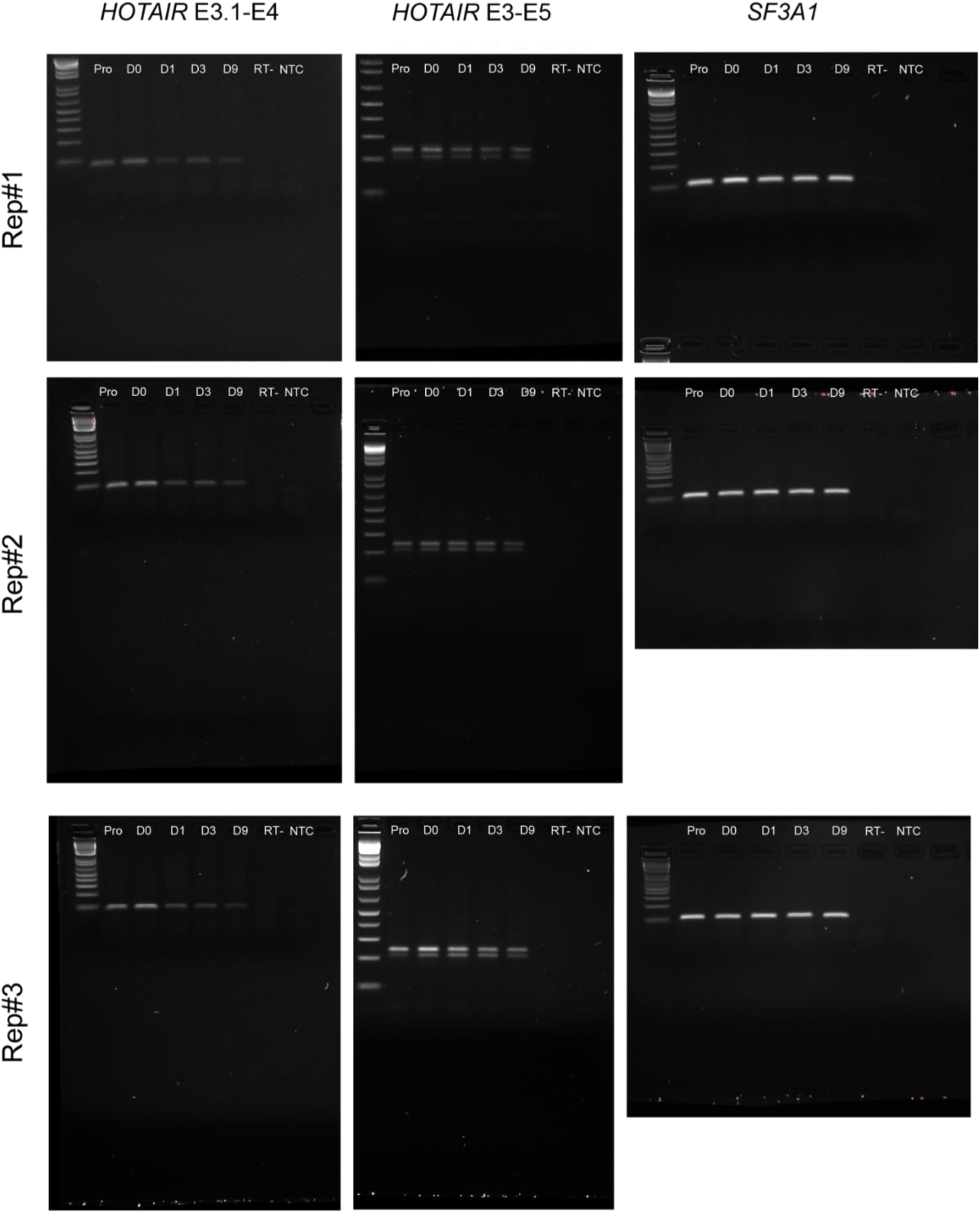
Uncropped gels for **Additional file 2, Fig. S2b**

**Figure S5.**
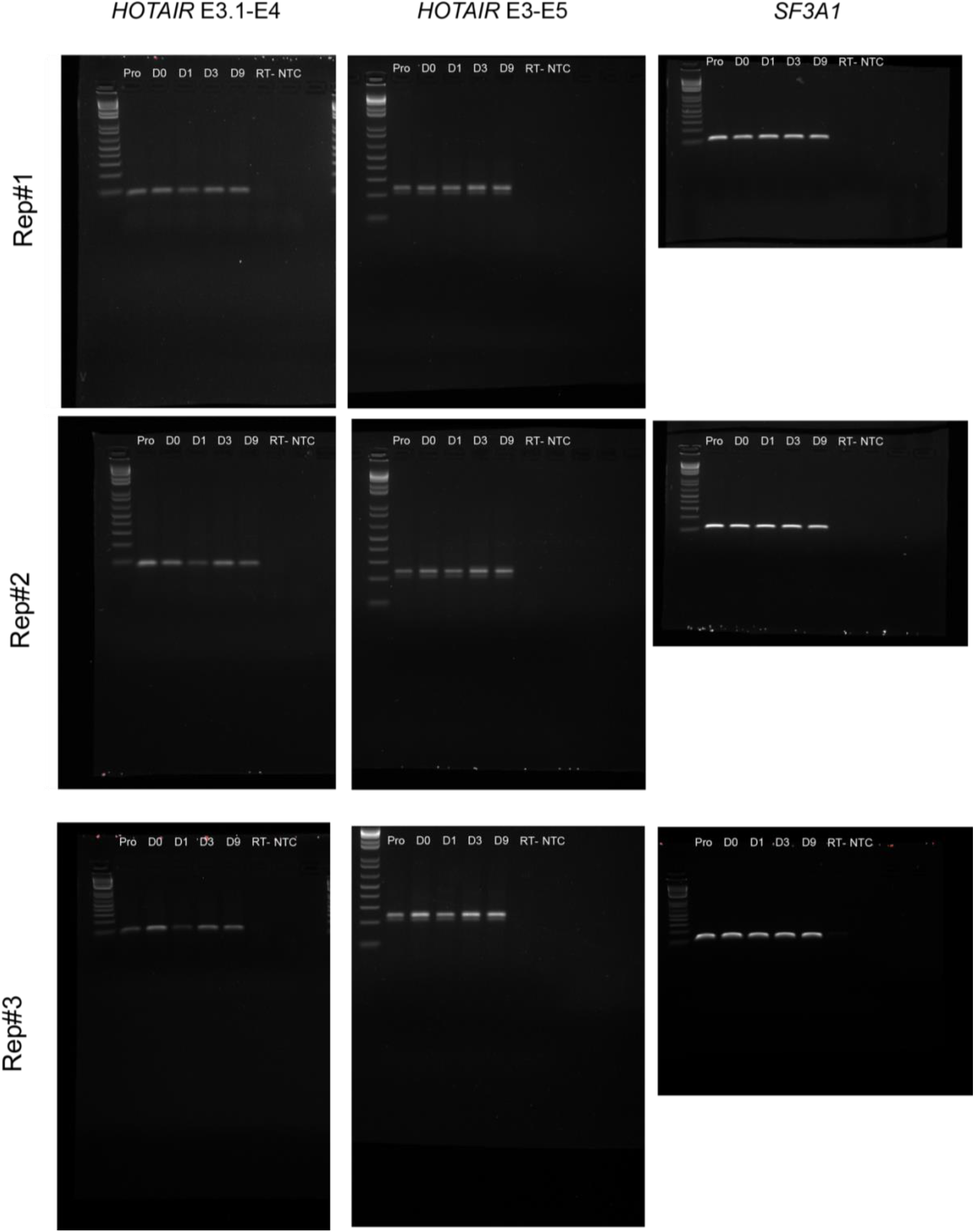
Uncropped gels for **Fig.4f** and **Additional file 2, Fig. S2a** right panel

**Figure S6.**
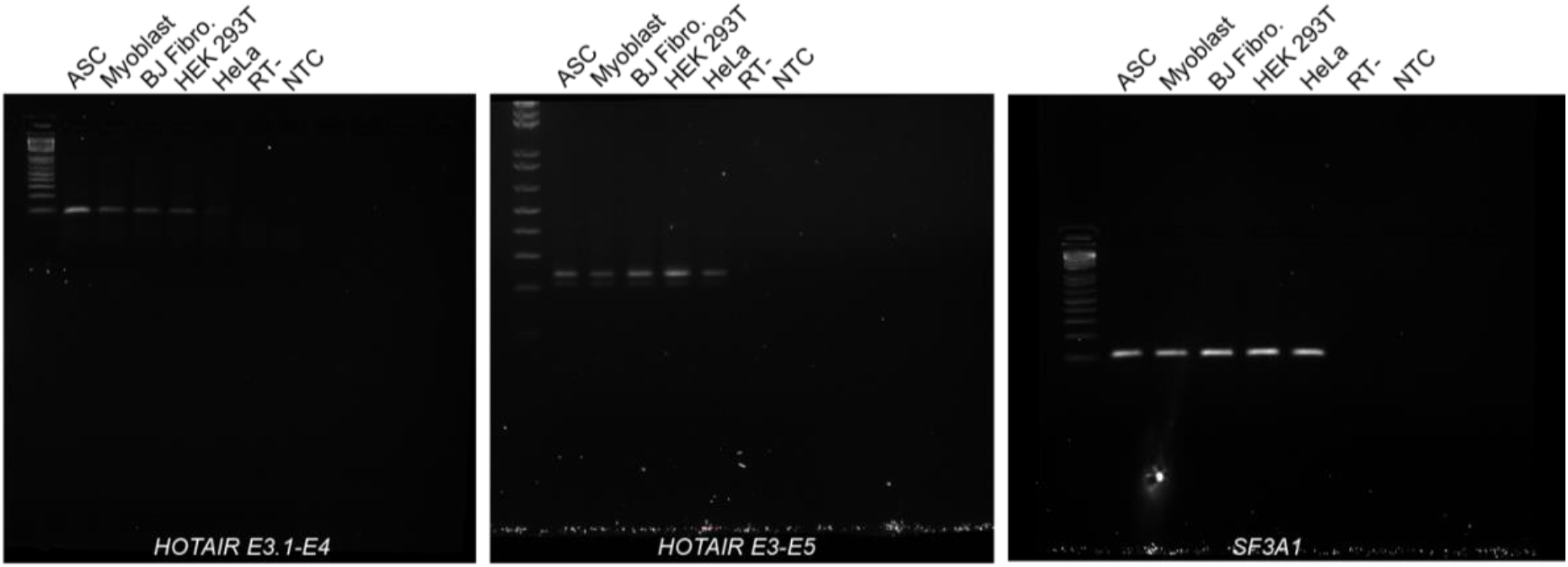
Uncropped gels for **Fig.5b**

**Figure S7.**
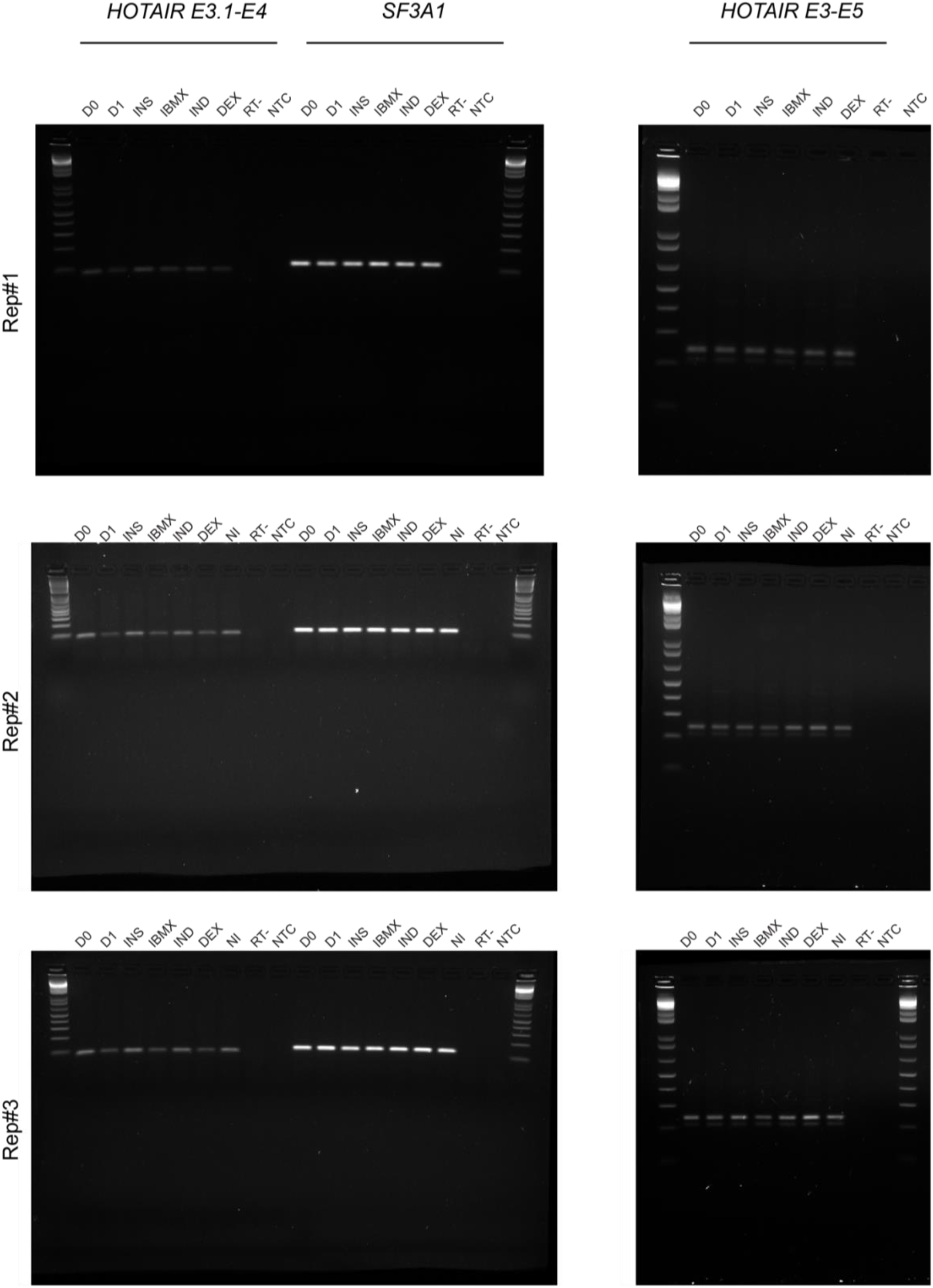
Uncropped gels for **Fig.7b** and **Additional file 2, Fig.S2c**

